# ERK1/2-dependent TSPO overactivation associates with the loss of mitophagy and mitochondrial respiration in ALS

**DOI:** 10.1101/2023.01.21.524879

**Authors:** Andrea Magrì, Cristiana Lucia Rita Lipari, Pierpaolo Risiglione, Stefania Zimbone, Francesca Guarino, Antonella Caccamo, Angela Messina

**Author notes:** Author to whom correspondence should be addressed: Angela Messina, Dept. Biological, Geological and Environmental Sciences, University of Catania, Via S. Sofia 64, 95123 Catania, Italy, Mail. These authors share the senior authorship (Antonella Caccamo, Angela Messina).

## Abstract

Mitochondrial dysfunction and the loss of mitophagy, aimed at recycling irreversibly damaged organelles, contribute to the onset of amyotrophic lateral sclerosis (ALS), a fatal neurodegenerative disease affecting spinal cord motor neurons. In this work, we showed that the reduction of mitochondrial respiration, exactly oxygen flows linked to ATP production and maximal capacity, correlates with the appearance of the most common ALS motor symptoms in a transgenic mice model expressing SOD1 G93A mutant. This is the result of the equal inhibition in the respiration linked to complex I and II of the electron transport chain, but not their protein levels. Since the overall mitochondrial mass was unvaried, we investigated the expression of the Translocator Protein (TSPO), a small mitochondrial protein whose overexpression was recently linked to the loss of mitophagy in a model of Parkinson’s disease. Here we clearly showed that levels of TSPO are significantly increased in ALS mice. Mechanistically, this increase is linked to the overactivation of ERK1/2 pathway and correlates with a decrease in the expression of the mitophagy-related marker, Atg12, indicating the occurrence of impairments in the activation of mitophagy. Overall, our work sets out TSPO as a key regulator of mitochondrial homeostasis in ALS.

## Introduction

Amyotrophic lateral sclerosis (ALS) is the most common, adult-onset disorder affecting the motor system, although extra-motor manifestations are emerging [1]. ALS triggers the progressive degeneration of upper and lower motor neurons (MNs) in the brain stem and spinal cord, leading to muscle weakness, atrophy, paralysis, and death within a few years from the appearance of the first symptoms [2–4]. Although ALS is generally sporadic (sALS), in a tenth of cases it is inherited with an autosomal dominant pattern, with more than 20 genes associated with familial ALS (fALS) forms [5]. Of these, mutations in the gene encoding the antioxidant enzyme Cu/Zn Superoxide Dismutase (SOD1) account for up to 20% of familial and about 5% of sporadic cases [6,7]. The mechanisms leading to the selective death of MNs in ALS are heterogeneous and not fully understood yet. Despite this, fALS and sALS are presumed to share the same pathological mechanisms, as they have similar clinical features [8].

Mitochondria are the energy supply stations of eukaryotic cells. Primarily, they produce ATP via oxidative phosphorylation, albeit they are essential for the maintenance of a variety of additional bioenergetic and biochemical pathways, including phospholipid biosynthesis, calcium homeostasis and cell survival [9,10]. Among cell types, neurons are the most susceptible to mitochondrial damage: they must last the lifetime of the organism and, having axons up to one meter long, require high energy demand for preserving their functions [11,12]. In light of these considerations, it is not surprising that mitochondria dysfunction is a key event in the onset of neurodegenerative diseases. Morphological and ultrastructural altered, swollen and vacuolated mitochondria were observed in many different ALS cases [13,14]. However, most of the literature information comes from SOD1 mutant models where impaired activity of the mitochondrial electron transport (ET) chain complex I and altered organelle dynamic were observed [15–18]. Remarkably, these alterations do not depend on the loss of dismutase activity of SOD1, but rather on the gain of toxic properties which prompts SOD1 misfolding and accumulation, and its partial re-localization from cytosol to mitochondria [19–24]. Within the organelle, mutated SOD1 presumably affects protein compositions, the integrity of the mitochondrial outer membrane (MOM), ATP/ADP trafficking mediated by VDAC1 and Bcl-2 activities [25–29].

To dispose of poorly functioning mitochondria, healthy MNs have a quality control system consisting in a specific form of autophagy, called mitophagy, aimed at recycling the components of irreparably damaged mitochondria [30]. Therefore, it is not surprising that the loss of mitophagy is rapidly emerging as a hallmark of neurodegenerative diseases. Initially, this aspect was investigated in Parkinson’s disease (PD), due to the direct involvement in disease pathogenesis of the E3 ubiquitin ligase Parkin and PTEN-induced kinase 1 (PINK1), both essential for the initiation of mitophagy process [31]. However, mutations in genes regulating mitophagy, such as those encoding optineurin and p62/sequestrosome-1, have been discovered also in ALS patients and linked to the disease onset [32,33]. In addition, alterations in this pathway were observed in models overexpressing mutated SOD1, TAR DNA-binding protein 43 (TDP-43) and FUS [34–36], all genes linked to ALS. More recently, a mitochondrial key regulator of mitophagy was identified in the Translocator Protein (TSPO) [37]. TSPO is an 18 kDa multi-drug binding protein located in the MOM found upregulated in PD patients [38]. It has been demonstrated that its expression is driven by the activation of MAPK/ERK pathway [39].

With the aim of further investigating this disease aspect, here we demonstrated that in transgenic mice expressing a G93A mutant form of human SOD1, impairments of motor abilities correlate with a significant reduction of mitochondrial respiration, precisely the oxygen consumption linked to ATP production and the maximal capacity, as the result of partial inhibition of respiration coupled to complexes I and/or II. This, however, is not due to a reduction in the mitochondrial mass but rather is the result of overactivation of ERK1/2 pathway, which induces TSPO overexpression via STAT3, in turn triggering a partial inhibition of mitophagy. Therefore, here we demonstrate that TSPO is a key regulator of mitochondrial quality control also in ALS.

## Results

### Congenic SOD1 G93A mice develop typical ALS phenotype

In this work, we used the commercially available congenic strain B6.Cg-Tg(SOD1*G93A)1Gur/J (here simply referred as transgenic mice). In comparison to the mostly used mixed B6SJL background carrying the same mutation, this strain shows an increased life span of about 20%, with the appearance of the overt symptoms at ∼20 weeks of age [40]. At this age, as expected, we noticed high expression of SOD1 in spinal cords of transgenic mice compared to wild type (Fig. 1A).

**Figure 1.**
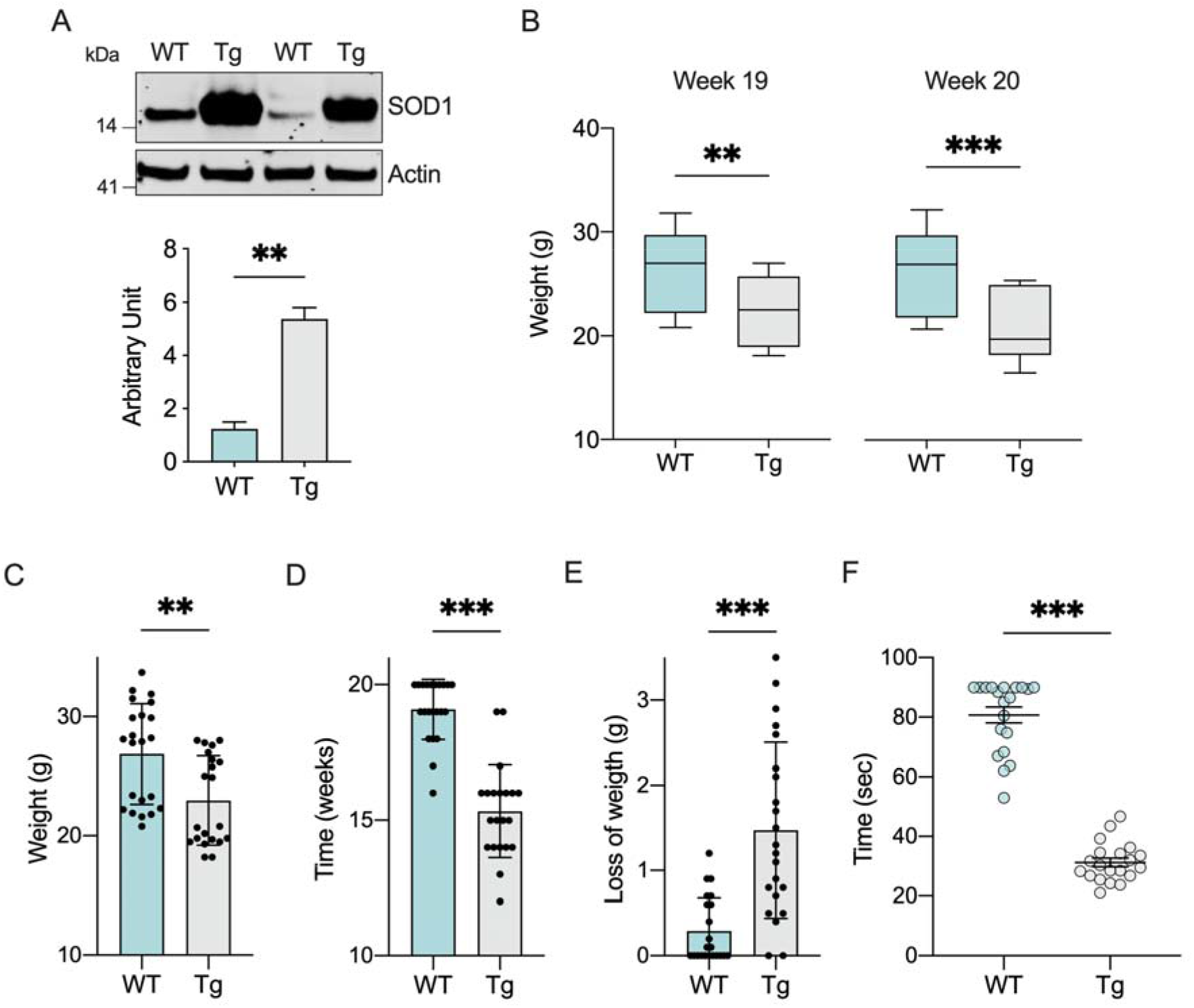
SOD1 G93A overexpression correlates with weight loss and motor impairment onset in transgenic mice. **A)** Representative Western blot of proteins extracted from spinal cord of wild type and transgenic mice and the relative quantitative analysis of SOD1. The loading control Actin was used for normalization. Data are expressed as mean ± SEM of n=2 independent measurements and statistically analyzed by Student t-test, with ** p<0.01. **B)** Analysis of body weight related to weeks 19 and 20. Data are expressed as median ± SEM of n=20 independent measurements and statistically analyzed by Student t-test, with ** p<0.01 and *** p<0.001. **C-E)** Maximum weight reached (C), age at the maximum weight (D) and the weight loss at the week 20 (E) relative to wild type and transgenic mice. Data are expressed as mean ± SD of n=20 independent measurements and analyzed by Student t-test, with ** p<0.01 and *** p<0.001. **F)** Rotarod test performed on accelerating rod at week 20. Data are expressed as mean ± SEM of n=20 independent measurements and analyzed by Student t-test, with *** p<0.001.

Female and male transgenic and wild type mice were further evaluated starting at 1 month of age for body weight changes (Suppl. Fig. 1). As shown in Fig. 1B, we observed a significant reduction of about 13.5% at week 19 in the body weight of transgenic mice (p=0.0043, n=20) and 18.2% at week 20 (p=0.0003, n=20). Particularly, transgenic mice reached a maximum weight value lower than wild type (22.9 ± 3.74 *vs*. 26.8 ± 4.22 g of control, p=0.0027, n=20, Fig. 1C) and at a significant younger age (15.3 ± 1.7 *vs*. 19.1 ± 1.1 weeks of control, p<0.001, n=20, Fig. 1D). Furthermore, the disease progression correlated with a significative weight loss in transgenic mice at the week 20 (1.54 ± 1 *vs*. 0.29 ± 0.38 gr of control, p<0.001, n=20, Fig. 1E).

To evaluate motor functions, rotarod test was performed. 19 weeks old transgenic and wild type mice were trained for two days on a rod at the constant speed; then, probe trials were conducted on day 3 on an accelerating rod (1 rpm/s, up to 40 rpm). In comparison to wild type, transgenic mice showed motor impairments already during the training (Suppl. Fig. 2), with a time spent on the rod of 74.8 ± 15.7 sec the first day and 75.6 ± 19.7 sec the second day (wild type performances were 88.6 ± 4.2, p<0.001, and 89.7 ± 1.3, p=0.0029, n=20, for day 1 and day 2, respectively). The difference between the two groups was exacerbated by the accelerating rod, where the average time spent on the rod was 30.7 ± 6.8 sec for transgenic and 81.2 ± 11.7 sec for wild type mice (p<0.001, n=20, Fig. 1F).

### Impaired mitochondrial respiration is observed in transgenic mice

To evaluate mitochondrial functionality, we investigated the oxygen consumption upon different conditions by using high-resolution respirometry on fresh spinal cord tissue homogenates. Fig. 2A shows a representative respirometric curve obtained from a wild type mouse alongside the specific Substrates-Uncoupled-Inhibitors Titration (SUIT) protocol used here (see Suppl. Tab 1 for raw data). Briefly, the non-phosphorylating respiration (LEAK state) was measured after the addition of the homogenate in the cuvette, in the presence of NADH-linked substrates (pyruvate, malate and glutamate) but not adenylates [41]. Then, respiration linked to the activation of ET complexes, i.e. the oxidative phosphorylation (OXPHOS state), was assayed in the presence of saturating concentration of ADP, before and after the addition of succinate. Maximal electron input to ET chain (ET capacity) was induced with uncoupler carbonyl cyanide 3-chlorophenylhydrazone (CCCP) titration, before and after the addition of rotenone. Finally, the residual respiration (ROX) was measured by the addition of antimycin.

**Figure 2.**
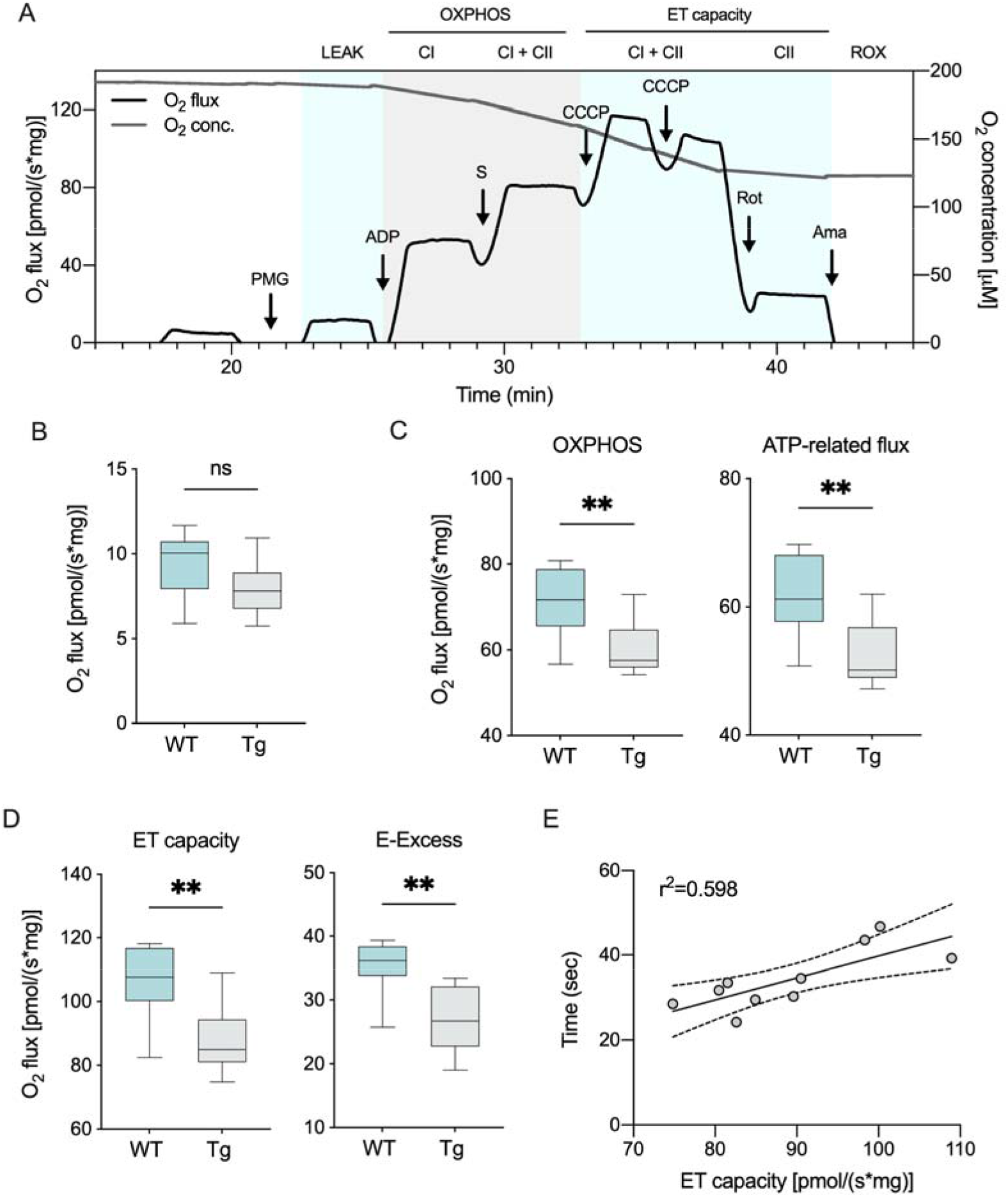
Mitochondrial respiration is impaired in transgenic mice. **A)** Representative trace of oxygen consumption achieved using homogenates from fresh spinal cord from wild type mice together with the protocol used. Precisely, the LEAK state was measured in the presence of pyruvate, malate and glutamate (PMG) but not adenylates. The addition of ADP, followed by succinate (S), activated OXPHOS respiration sustained by complex I or complex I and II, respectively. The maximal capacity of ET chain was achieved by CCCP titration. The ET capacity sustained exclusively by complex II was measured after inhibition of complex I with rotenone (Rot). Finally, the ROX was measured after a complete inhibition of ET chain with antimycin A (Ama). **B-D)** Comparative analysis of oxygen consumption in wild type and transgenic mice spinal cords’ homogenates relative to LEAK (B), OXPHOS, total and ATP-linked flow (C), maximal ET capacity and E-excess (D). Data are expressed as pmol/s of oxygen per mg of tissue and as means ± SEM of n=10 independent measurements. Data were statistically analyzed by Student t-test, with ** p<0.01. **E)** Correlation analysis of ET capacity with the loss of motor activities calculated as linear regression (r^2^=0.598, p=0.0087). The dotted lines represent 95% confidence intervals.

While no differences were observed between the groups for the LEAK respiration (Fig. 2B), in comparison to wild type, transgenic mice show a significant reduction of the OXPHOS respiration: upon stimulation of the ET chain with externally added substrates and ADP, oxygen consumption was reduced of about 15% (p=0.0056, n=10, Fig. 2C); accordingly, the OXPHOS flux coupled to ADP phosphorylation, the so-called net flux, decreased in a proportional manner in transgenic mice (p=0.004, n=10, Fig. 2C). Similarly, ALS pathology affects both maximal ET capacity of mitochondria and the E-Excess, a respiratory reserve used by mitochondria in stress conditions. As showed in Fig. 2D, both parameters were significantly reduced in transgenic mice (respectively, p=0.0023 and p=0.0012, n=10). Furthermore, the reduction of maximal ET capacity in transgenic mice positively correlates with the impairment in motor capacity, i.e., the time spent on the accelerating rod (Fig. 2E, r^2^=0.589, p=0.0087). Overall, our data indicate that reduction of mitochondrial respiration occurs as the phenotypical manifestations of ALS appear.

### Complex I and II linked respiration, but not their expression or mitochondrial mass, is reduced in transgenic mice

Based on previous results, we queried whether the impairment of ET chain functioning observed in transgenic mice was due to a reduction in the activity and/or protein expression of the ET complexes that in our set-up activate the chain, as well as mitochondrial mass. The SUIT protocol in Fig. 2A, indeed, allows the measurement of respiration driven by complex I. As schematized in Fig. 3A, this was achieved upon stimulation with the NADH-linked substrates and saturating concentration of ADP, but not succinate that specifically activates complex II: in this configuration, electrons flow to complex III exclusively from complex I via the Q-junction. As reported in the histogram of Fig. 3A, a significant reduction of complex I linked oxygen flux was observed in transgenic mice compared to wild type (− 20%, p=0.0019, n=10). This data is in accordance previous reports in cell lines and other mouse models indicating a specific impairment of complex I activity [28,42]. To analyze if the protein levels of complex I were altered, we measured the expression of NADH ubiquinone oxidoreductase core, NDUFV1. As reported in Fig. 3B, no differences were detected between groups (p=0.97, n=4). Similar results were obtained for the protein levels of Tom20, a widely marker used to assay mitochondrial mass (p=0.69, n=4).

**Figure 3.**
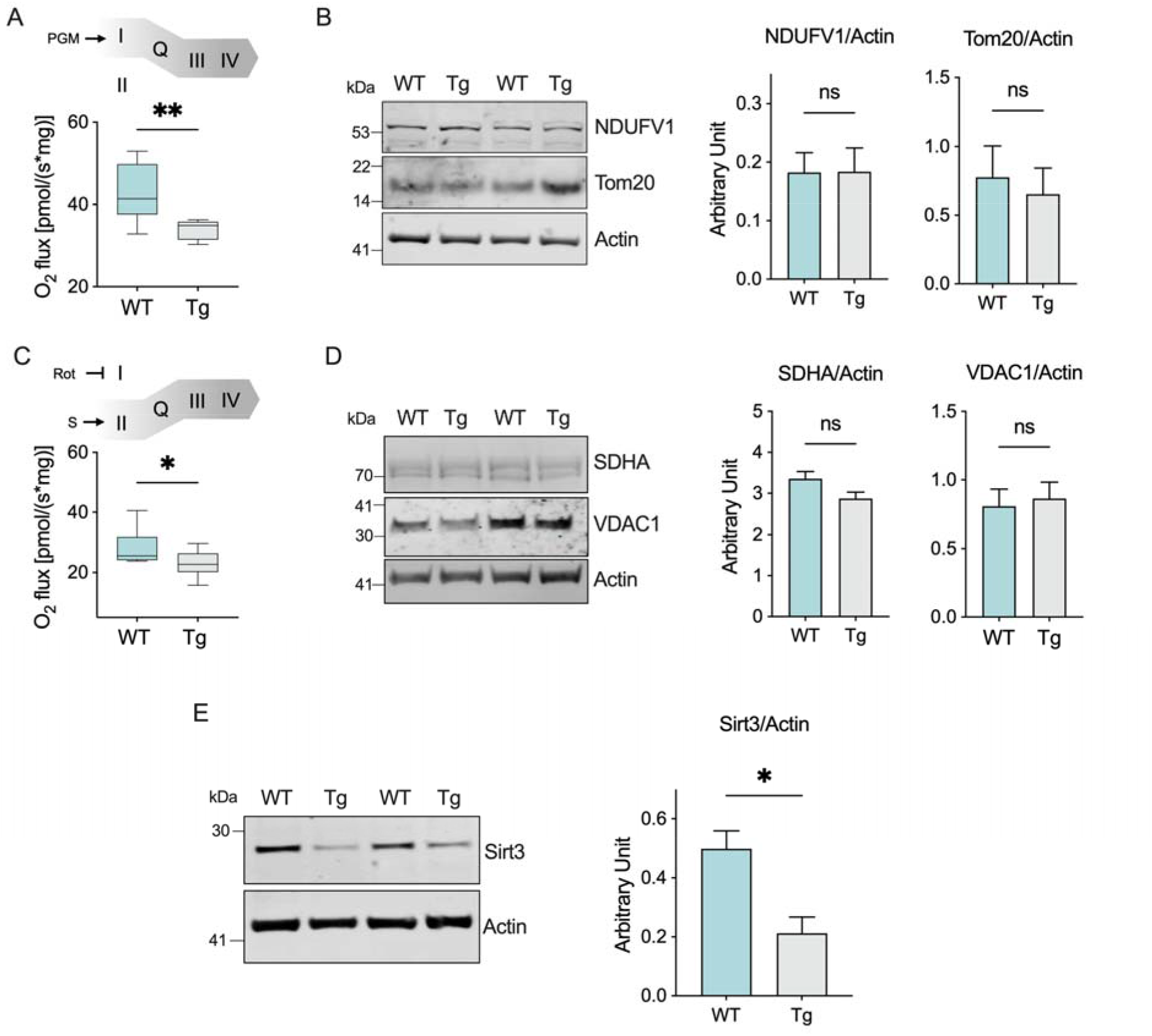
Transgenic mice show a reduced contribution in complexes I and II of the ET chain functioning but not in the mitochondrial mass. **A)** Schematic representation of the experiment rationale and comparative analysis of oxygen consumption in wild type and transgenic mice spinal cords’ homogenates relative to the OXPHOS sustained by complex I. **B)** Western blot of proteins extracted from spinal cord of wild type and transgenic mice and the relative quantitative analysis of NDUFV1 and Tom20. **C)** Schematic representation of the experiment rationale and comparative analysis of oxygen consumption in our mice groups relative to the ET capacity sustained by complex II. **D)** Western blot and the relative quantitative analysis of SDHA and VDAC1 in our samples. **E)** Western blot and the relative quantitative analysis of Sirt3 in our samples. Data in (A) and (C) are expressed as pmol/s per mg of tissue and as mean ± SEM of n=10 independent measurements. Data were statistically analyzed by Student t-test, with * p<0.05 and ** p<0.01. Western blot quantifications in (B), (D) and (E) were performed by normalizing the protein of interest to Actin, here used as loading control. Data are expressed as mean ± SEM of n=4 independent measurement and statistically analyzed by Student t-test, with * p<0.05; *n*.*s*. not significant.

Next, the respiration coupled to complex II (succinate dehydrogenase, SDH) was investigated as the specific contribution to the ET capacity in the presence of succinate and after the complete inhibition of complex I with rotenone (Fig. 3C). Again, complex II linked respiration was significantly reduced in transgenic mice in comparison to wild type (− 20%, p=0.03 *vs*. wild type, n=10, Fig. 3C). Similarly to what observed previously, we did not notice any variations in the protein levels of SDHA, the main hydrophobic subunit of SDH, or VDAC1, the most abundant protein of the MOM (p=0.1 and p=0.75 respectively, n=4, Fig. 3D). Being directly involved in the modulation of the overall mitochondrial metabolic activities and acting as a physiological regulator of SDH functioning [43], we then assayed the expression levels of the NAD-dependent deacetylase sirtuin-3 (Sirt3). As reported in Fig. 3E, Sirt3 was significantly downregulated in transgenic mice compared to wild type (p=0.012, n=4).

### High expression of TSPO, activated by ERK1/2, inhibits mitophagy in transgenic mice

Despite a general impairment of the ET chain activity, we did not detect any variation in the expression levels of proteins used as marker of mitochondrial mass and respiration complexes. This suggest that damaged mitochondria may accumulate in the MNs of transgenic animals. To explore this theory, we looked at TSPO, a small protein located in the MOM with a well-characterized role in the regulation of mitochondrial metabolism and a known marker of neuroinflammation and microglia activation [44]. Using a cellular model of PD, it was recently demonstrated that its overexpression correlates with a significant inhibition of mitophagy [37]. Therefore, we measured the levels of TSPO expression in total lysates of spinal cords by western blot. As reported in Fig. 4A, the protein levels of TSPO were about three times higher in transgenic mice compared to wild type (p=0.0004, n=4).

**Figure 4.**
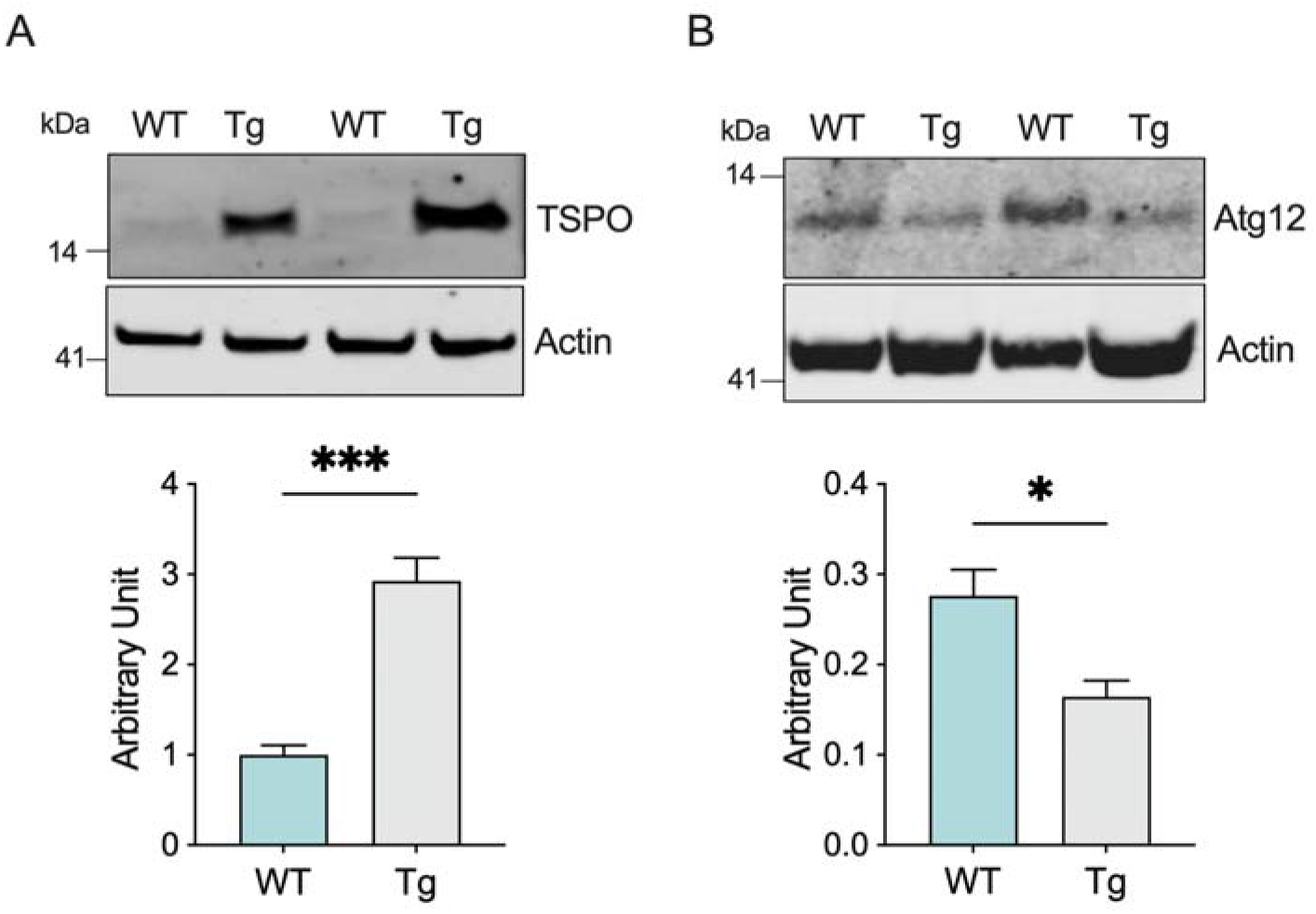
Expression of TSPO and Atg12 are inversely correlated in transgenic mice. Western blot of proteins extracted from spinal cord of wild type and transgenic mice and the relative quantitative analysis of TSPO **(A)** and Atg12 **(B)**. Western blot quantifications were performed by normalizing the protein of interest to Actin, here used as loading control. Data in histograms are expressed as mean ± SEM of n=4 independent measurement and statistically analyzed by Student t-test, with * p < 0.05 and *** p < 0.001.

To assess whether the increase in TSPO expression affected mitophagy, we measured the protein levels of the autophagy marker Atg12, which represents one of the fundamental components of the autophagosome [45]. As displayed in Fig. 4B, Atg12 levels were significantly reduced in transgenic mice: precisely, densitometric analysis revealed a reduction of about 40% in comparison to wild type (p=0.016, n=4).

It is known that TSPO expression is regulated by the MAPK/ERK pathway via STAT3 [39]. Furthermore, it has been hypothesized that overactivation of ERK1/2 signaling cascade could play a crucial role in the development of ALS [46]. In light of these considerations, we assessed the levels of expression and phosphorylation of the proteins involved in this pathway. As showed in Fig. 5A, while the overall protein levels of ERK1 and ERK2 remained substantially unchanged, a significant increasement of the protein phosphorylation was observed exclusively in transgenic mice. Densitometric quantification indicated that levels of phosphorylated ERK1 and ERK2 were more than doubled in transgenic compared to wild type mice (Fig. 5B-C, p<0.0001, n=4). Accordingly, the downstream effector STAT3 was similarly activated: as displayed in Fig. 5A and 5D, the levels of STAT3 were similar between groups albeit the phosphorylated form was detected exclusively in transgenic mice (p=0.0002, n=4).

**Figure 5.**
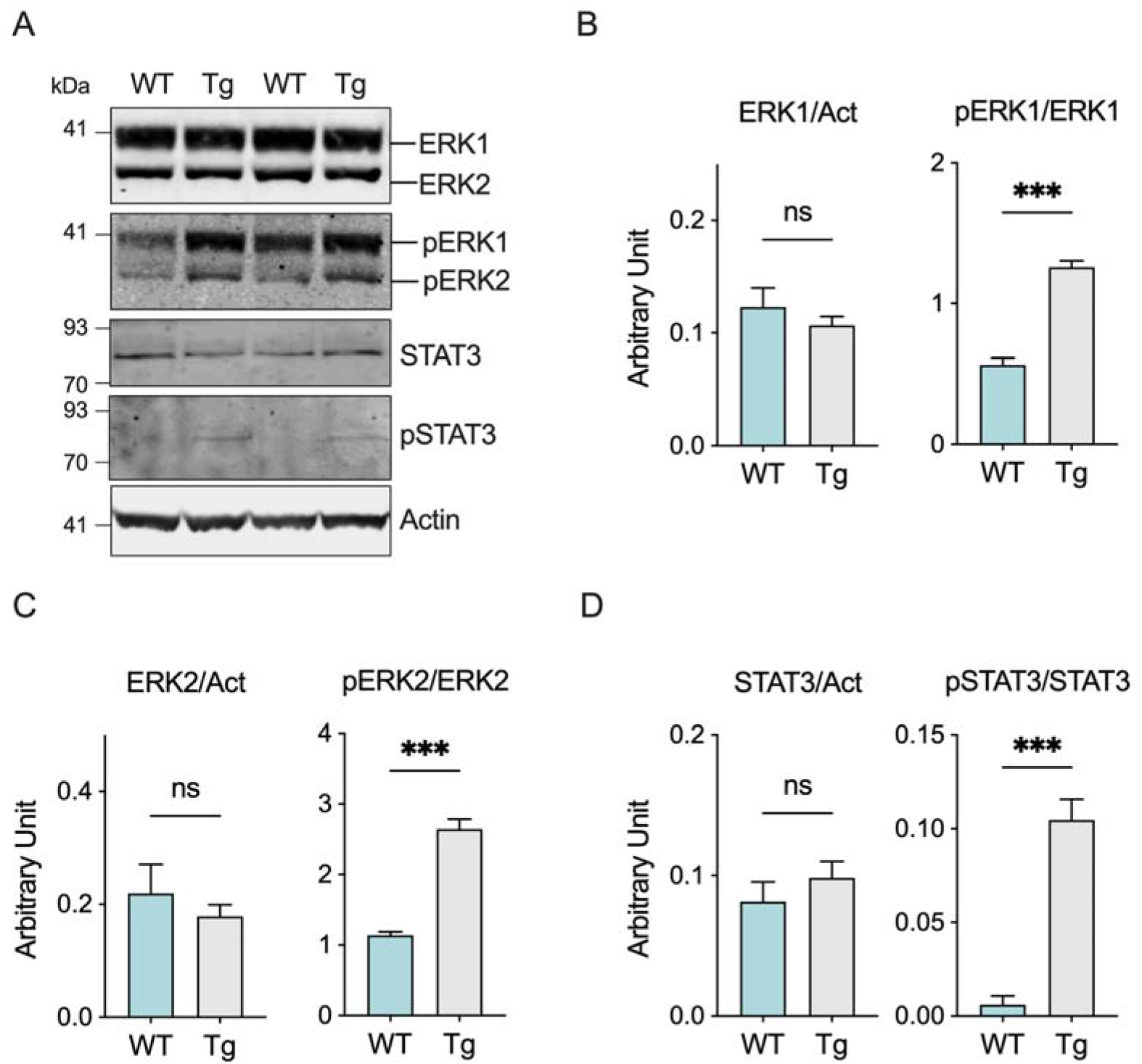
ERK1/2 signaling is activated in ALS transgenic mice. **A)** Western blot of proteins extracted from spinal cord of wild type and transgenic mice relative to total and phosphorylated ERK1/2 and STAT3. **B-D)** Quantitative analysis of total and phosphorylated ERK1 (B), ERK2 (C) and STAT3 (D). Western blot quantifications of total proteins were performed by normalizing the protein of interest to Actin, while quantification of phosphorylated proteins were obtained using the relative total protein for normalization. Data are expressed as mean ± SEM of n=4 independent measurement and statistically analyzed by Student t-test, with *** p < 0.001; *n*.*s*. not significant.

These data clearly indicate that ERK1/2-dependend TSPO overactivation correlates with the inhibition of autophagosome formation, thus with an alteration of the mitophagy process.

## Discussion

Although the large heterogeneity of the genetic and sporadic ALS, mitochondrial dysfunction characterizes all forms of the disease. Metabolic impairments arise in the early stages of the disease and rapidly culminate into morphological, ultrastructural and functional abnormalities [47,48]. Furthermore, as emerges from recent literature, MNs affected by the disease show dysregulation of autophagy, which hinders the recycling of dysfunctional organelles and further aggravates the MNs function. Indeed, defects in autophagy are also responsible for the accumulation of protein aggregates and reactive oxygen species (ROS) as well as increased inflammation, all conditions that characterize ALS and other neurodegenerative diseases [49].

To try to understand molecular mechanisms behind this intricate puzzle, here we studied the mitochondrial dysfunction in a murine model of ALS. We found that: i) the onset of motor manifestation correlates with a general decrease in oxygen consumption but not in mitochondrial mass; ii) high level of TSPO protein expression found in the spinal cord correlates with an inhibition of mitophagy, similarly to what previously observed in PD [37].

In this work, we used the SOD1 G93A mouse model of ALS that, as expected, develops the typical ALS motor symptoms (Fig. 1). Around 20 weeks of age, spinal cords’ mitochondria show a strong impairment of their functionality as demonstrated by the reduced oxygen consumption rates observed in different respiratory states. Precisely, we confirmed a significant reduction of OXPHOS-linked respiration (Fig. 2C) and the maximal capacity of the ET system (Fig. 2D). Overall, reduction of these oxygen flows has at least two important consequences. First, the ATP-related OXPHOS flux is proportionally reduced (Fig. 2C), confirming previous data about ATP production, a key event that triggers mitochondrial dysfunction [26]. Second, the so-called E-Excess is dramatically compromised (Fig. 2D). Particularly, excess capacity is a crucial reserve to which mitochondria may rely on in cases of increase in energy demands or stress conditions [41]. In fact, the reduction of respiratory reserves are considered prodromal to mitochondrial dysfunction in many pathologies [50], as we recently observed in two different models of PD [51–53]. Thus, the concomitant reduction of ATP availability and E-Excess makes mitochondria more susceptible to further toxic insults.

Notably, reduction of respiration coupled to complex I and II, here calculated as the specific contribution of each complex in the activation of the ET chain and analyzed in real time upon the addition of specific substrates (Fig. 3A and C), is not related to variation in the protein levels of specific complexes subunits, nor to changes in mitochondrial mass, as demonstrated by Western blot in Fig. 3B and D. Certainly, inhibition of complex I in ALS is known since 1998 [54] and is considered a consequence of the combination of different factors, including the limited availability of NADH-linked substrates due to the deposition of SOD1 mutants on the MOM [24,26,55,56]. Additionally, we found a significative reduction of Sirt3 expression in transgenic mice (Fig. 3E) that can explain the deficiency in the ET chain activity. Sirt3 is the major mitochondrial deacetylase that controls enzymes activity (including those of the ET chains), the organelle integrity and ROS homeostasis [57,58]. In particular, SDH acts as a binding partner and substrate for the deacetylase activity of Sirt3: therefore, a reduction in Sirt3 expression may result in a decrease of SDH enzymatic activity [43]. Remarkably, downregulation of Sirt3 was recently observed in MNs derived from iPSC of both sporadic and familial ALS patients where it correlated with a reduction in mitochondrial respiration [59]. In lights of these considerations, it is conceivable that reduced protein levels of Sirt3 contribute to the impairment of ET activity.

In parallel with mitochondrial dysfunction, we noticed significant changes in TSPO expression. Physiologically, TSPO participates in a variety of mitochondrial and cellular functions, such as cholesterol import into the organelle, synthesis of heme and steroid hormones, anion transport, cell proliferation and apoptosis [60–62]. Moreover, TSPO has been widely used in the last decades as biomarker for brain imaging because was found to increase upon injury, inflammation and the onset of neurodegenerative disorders [44,63–[66]. Interestingly, only recently the Campanella’s group linked TSPO overexpression to loss of mitophagy in a cellular model of PD [37].

We observed a similar condition in our ALS model. Exactly, our results show that, in healthy spinal cord, TSPO expression is very low, consistent with data from a previous report [67], whereas in ALS its level is about three time higher (Fig. 4A). This correlates with a significant reduction of the autophagy related protein Atg12 (Fig. 4B). Notably, Atg12 is, among Atg proteins, the one directly related to mitophagy: indeed, its overexpression exerts a protective role against mitochondrial stress insults by selectively stimulating activation of mitochondrial quality control systems [68]. Consistent with increased levels of TSPO, we noticed a significant activation of ERK1/2 signaling and STAT3 (Fig. 5), both essential for TSPO transcription [39].

ERK1/2 signaling represents one of the most important intracellular cascades for extracellular stimuli. In the nervous system, activation of ERK1/2 in response to growth factors, chemokines, cytokines and oxidative stress exerts a protective role, being involved in cell proliferation, survival and differentiation [69]. On the other side, it has been proposed that the excessive ERK phosphorylation causes neuronal abnormalities in ALS [46]. For instance, in an ALS model overexpressing TDP-43, ERK1/2 hyperactivation correlates with the accumulation of TDP-43 positive aggregates [70] and with an increase in the phosphorylation of the ERK1/2 downstream effector, p90SRK [71]. Similarly, in SOD1 G93A mice an increase in ERK1/2 phosphorylation was observed in different areas of central nervous system as the pathology progresses [72]. Remarkably, ERK1/2 activation has a direct effect on mitochondrial functioning: when induced by oxidants, ERK1/2 triggers impairment of mitochondrial respiration, ATP production and substrate oxidation acting on complex I function [73]. Taken together, these data that confirm the results found in this work. Thus, on one hand, ERK1/2 activation promotes TSPO overexpression via STAT3, triggering in turn mitophagy impairment; on the other hand, ERK1/2 could directly affect the ET chain function, that is already compromised by the accumulation of SOD1 mutants on the MOM and/or the reduced Sirt3 levels (see Fig. 6 for a schematization of the proposed model). Overall, these data clearly show for the first time a possible correlation between the overactivation of TSPO pathway with alterations in mitochondrial respiration and mitophagy in ALS.

**Figure 6.**
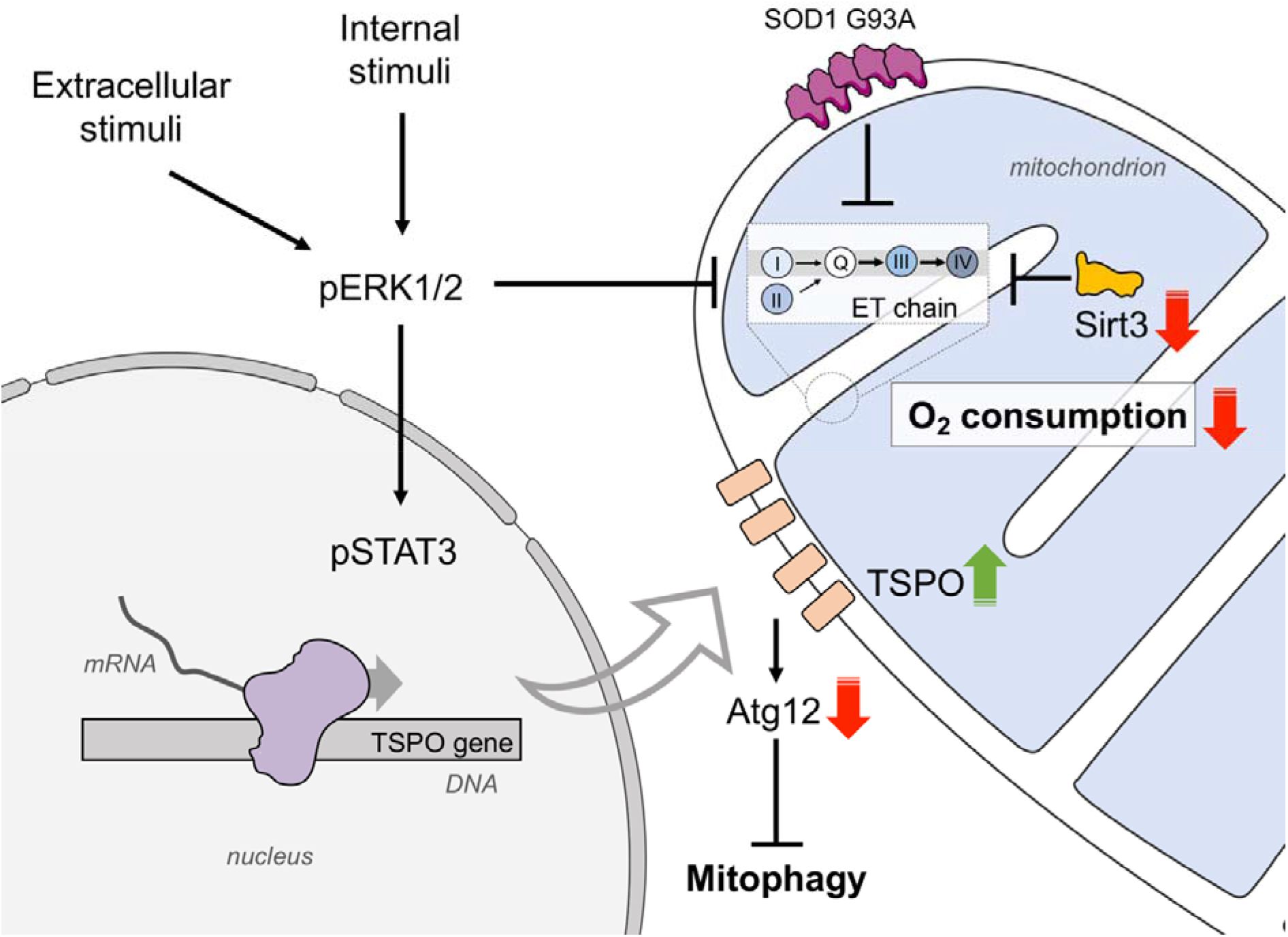
Proposed model on the role of ERK1/2 and TSPO in ALS and their involvement in mitochondrial dysfunction. The activation of ERK1/2 signaling in ALS is the sum of both external and internal stimuli and promotes TSPO overexpression via STAT3. It has been recently demonstrated that high levels of TSPO prevent the activation of mitophagy. At the same time, ERK1/2 contributes to the reduction of ET chain functioning. Notably, enzymatic activity of ET chain is already compromised in ALS motor neurons by a sum of converging factors, including the accumulation of SOD1 G93A in the mitochondria and the lower levels of Sirt3.

The pathogenesis of ALS has not yet been fully elucidated, and the most accepted hypotheses attribute considerable significance to excitotoxicity, neuroinflammation, oxidative stress, and mitochondrial dysfunction. In particular, studies on mitophagy and its correlation with inflammation have become of special interest. Indeed, it is now well understood that improving mitochondrial autophagy may represent an opportunity to restore mitochondrial homeostasis, thus offering important insights into the development of potential treatments for ALS.

## Materials and Methods

### Animals

Transgenic mice B6.Cg-Tg(SOD1*G93A)1Gur/J (JAX ref. no. 004435), expressing high copy number of human SOD1 G93A mutant [74], and C57BL6/J wild type mice were purchased from The Jackson Laboratory (Bar Harbor, ME, USA). The colony was maintained by breeding male hemizygous carriers to wild-type females in order to obtain transgenic and wild-type mice. A sample size of 40 mice, 20 wild-type and 20 transgenic, with an equal number of males and females, at the age-matched of 20 weeks-old was used in this study. Sample size was estimated on the basis of information from previous studies from our and independent groups [42,75,76]. Mice were housed 4–5 to cage, kept on 12 h light/dark cycle and were given *ad libitum* access to food and water.

All experimental procedures were carried out according to the Italian Guidelines for Animal Care (D.L. 116/92 and 26/2014) and in compliance with the European Communities Council Directives (2010/63/EU). Protocols were approved by the Ethical Committee for animal experimentation at the University of Catania (OPBA, project ref. no. 334). All measures were adequately taken to minimize the number of animals.

### Genotypization

Genomic DNA was extracted from small tail biopsies by overnight digestion at 55°C in lysis buffer (0.1 M Tris-HCl, pH 7.5, 5 mM EDTA, 0.2 M NaCl, 0.2% SDS) supplemented with 0.4 mg/ml of proteinase K (Sigma-Aldrich, St. Luis, MO, USA). DNA was then isolated by isopropanol-based separation and resuspended in TE buffer (10 mM Tris-HCl, pH 7.5, 1 mM EDTA, 10 mM NaCl).

Transgenic mice were identified by PCR according to manufacturer protocol. The couple of primers 5’-CAT CAG CCC TAA TCC ATC TGA-3’ and 5’-CGC GAC TAA CAA TCA AAG TGA-3’ was used to amplify a 236 bp product from exon 4 of the hSOD1 gene within the transgene construct. The couple of primers 5’-CTA GGC CAC AGA ATT GAA AGA TCT-3’ and 5’-GTA GGT GGA AAT TCT AGC ATC ATC C-3’ was used to amplify a 324 bp product of the endogenous interleukin 2 gene, here used as DNA positive/internal control.

### Phenotypical analysis and rotarod test

To evaluate phenotypical changes, mice were weighed weekly starting at 1 month of age. Body weight loss was calculated as the difference between the maximum weight reached and the weight at the week 20. To evaluate motor functions, rotarod test was performed as in [76]. Briefly, mice were trained for 90 sec (4 trials per day, for 2 consecutive days) on a rod at a constant speed of 15 rpm. Then, 4 probe trials of 90 sec each were conducted on day 3 on an accelerating rod (1 rpm/s, up to 40 rpm). The entire set of animals was used.

### Spinal cord extraction and tissue homogenate preparation

Animals were sacrificed by CO_2_ asphyxiation at 20 weeks of age and spinal cords were rapidly removed. To preserve the mitochondrial functions and membranes integrity, tissues were kept in ice-cold BIOPS buffer (10 mM Ca EGTA, 20 mM imidazole, 20 mM taurine, 50 mM K-MES, 0.5 mM DTT, 6.56 mM MgCl_2_, 5.77 mM ATP, 15 mM phosphocreatine, Sigma-Aldrich) [77]. Alternatively, spinal cords were frozen in liquid nitrogen and stored at -80°C for further use.

Fresh spinal cords were gently homogenate within one hour from extraction by using a Potter Elvehjem tissue homogenizer in mitochondrial respiration buffer Mir06 (Oroboros Instruments, Innsbruck, Austria) and immediately used for respirometric analysis.

### High-resolution respirometry

The oxygen consumption of mitochondria was assayed in fresh spinal cord homogenates from transgenic or wild type mice by high-resolution respirometry, using the double-chamber system O2k FluoRespirometer (Oroboros Instruments). Tissue homogenization protocol was tested for mitochondrial membranes integrity by cytochrome c (cyt c) assay [77,78]. After the stimulation of respiration, 10 μM of cyt c was added to the cuvette and any eventual run showing significant a increase in oxygen consumption, due to the cyt c entry through damaged membranes, was excluded from the analysis.

A specific SUIT protocol aimed at investigating the main respiratory states was adapted from [77]. A volume of homogenate equivalent to 2 mg of the original tissue was added to the cuvette and the LEAK state was monitored in the presence of 10 mM pyruvate, 2 mM malate and 10 mM glutamate. The subsequent addition of saturating concentration of ADP (5 mM) allowed to measure OXPHOS respiration exclusively driven by complex I. Complex II was next activated by the addition of 10 mM succinate, measuring the total OXPHOS respiration. To assay the maximal ET capacity a titration with 0.5 μM of the uncoupler CCCP was performed. Finally, complex I was selectively inhibited with the addition of 2 μM rotenone to achieve ET capacity linked to complex II. With the addition of 2.5 μM antimycin, ROX was measured.

All the experiments were performed in Mir06 (Oroboros Instrument) at 37°C under constant stirring (750 rpm). All chemicals were purchased by Sigma-Aldrich. A set of animals of n=10 per experimental group, with an equal number of males and females, was employed for respirometric assays.

### Respirometric data analysis

Instrumental and chemical background fluxes were calibrated as a function of the oxygen concentration using DatLab software (version 7.4.0.1, Oroboros Instruments). Rate of oxygen consumption corresponding to LEAK, OXPHOS and ET capacity was expressed as pmol/s per milligram of tissue, and corrected for the ROX.

The ATP-related oxygen flux was calculated by normalizing the total OXPHOS flux for the LEAK respiration; the E-Excess was calculated as the difference between ET capacity and the OXPHOS respiration achieved in the presence of all substrates [51,79]. The Pearson correlation coefficient r was calculated for transgenic mice between maximal ET capacity value and the rotarod test output (i.e., the time spent on the rod) by using Prism software (version 9, GraphPad Inc., San Diego, CA, USA). A value of p<0.05was taken as significant.

### Western blotting

Spinal cord were homogenized in T-PER buffer (ThermoFisher) supplemented with cOmplete Protease Inhibitor Cocktail (Roche, Basel, Switzerland) and Halt Phosphatase Inhibitor Cocktail (ThermoFisher) to preserve protein integrity and post-translational modifications. Total protein lysates were quantified by Pierce BCA Protein Assay Kit (ThermoFisher) and added to NuPAGE LDS sample buffer, supplemented with sample reducing agent (ThermoFisher). Approximately 30 μg of proteins/sample were separated on NuPAGE Bis-Tris polyacrylamide gels (ThermoFisher) at 150 V for 50 mins. Proteins were then transferred to nitrocellulose membranes (GE Healthcare, Boston, MA, USA) using the semi-dry system PerfectBlue Electro Blotter (Peqlab, Germany), and the transfer was confirmed by using Ponceau S staining. Membranes were blocked in 5% BSA or not fat milk in PBS with 0.1% Tween-20. Full or portions of membranes were incubated overnight at 4 °C with primary antibodies against SOD1 (Abcam, Cambridge, UK, ref. no. ab16831, 1:1000), NDUFV1 (Immunological Sciences, ref. no. AB-83826, 1:500), Tom20 (Abcam, ref. no. ab186735, 1:500), SDHA (Abcam, ref. no. ab137040, 1:500), VDAC1 (Abcam, ref. no. ab14734, 1:1000), Sirt3 (Immunological Sciences, ref. no. AB-84353, 1:500), TSPO (Immunological Sciences, ref. no. AB-84352, 1:500), ERK1/2 (Cell Signaling, Danvers, MA, USA, ref. no. 4695, 1:2000), phospho-ERK1/2 (Cell Signaling, ref. no. 4370, 1:1000), STAT3 (Cell Signaling, ref. no. 4904,1:500), phospho-STAT3 (Cell Signaling, ref. no. 9145, 1:500), Atg12 (Cell Signaling, ref. no. 4180, 1:1000), β-Actin (Cell Signaling, ref. no. 3700, 1:2000). Then, membranes were incubated with IRDye conjugated secondary antibodies (LI-COR Biosciences, Lincoln, NE, USA, 1:25.000). Signals were detected using Odissey Imaging System (LI-COR Biosciences). Band quantification was performed by densitometric analysis using Image Studio Lite software (LI-COR Biosciences). A set of animals of n=4 per experimental group, with an equal number of males and females, was employed for respirometric assays.

### Statistical analysis

Data were expressed as a mean or mean ± SEM or SD and statistically analyzed by 2-way ANOVA followed by Sidak’s multiple comparison test or unpaired Student t-test using Prism software (version 9, GraphPad Inc.). The values of p<0.05, p<0.01 and p<0.001 were taken as significant.

## Acknowledgements

The authors are grateful to Salvatore Oddo (University of Messina) and Vito De Pinto (University of Catania) for their valuable support and for the critical discussion of the data. Authors also acknowledge Fondi di Ateneo 2020–2022, Università di Catania, linea Open Access, and Attraction and International Mobility (AIM) program, Linea 1, Salute.

## Availability of data and material

All data generated or analyzed during this study are included in this published article and its supplementary information files.

## Ethics approval and consent to participate

Not applicable

## Authors’ contributions

A Magrì performed behavior and respirometric experiments, analyzed and interpreted the data, draw pictures, wrote the original draft and reviewed the manuscript. CLRL participated in behavior experiments and performed of western blot. PR participated in respirometric and western blot experiments, and analyzed the data. SZ performed mouse breeding, genotypization and samples collection, and participated in behavior experiments. FG analyzed the data and reviewed the manuscript. AC designed the study, supervised all animal experiments, analyzed and interpreted the data, contributed to original draft preparation and reviewed the manuscript. A Messina conceptualized, designed and supervised the study, analyzed and interpreted the data, contributed to original draft preparation, reviewed and finalized the final version of the manuscript.

