## Supplementary Materials for "ERK1/2-dependent TSPO overactivation associates with the loss of mitophagy and mitochondrial respiration in ALS"

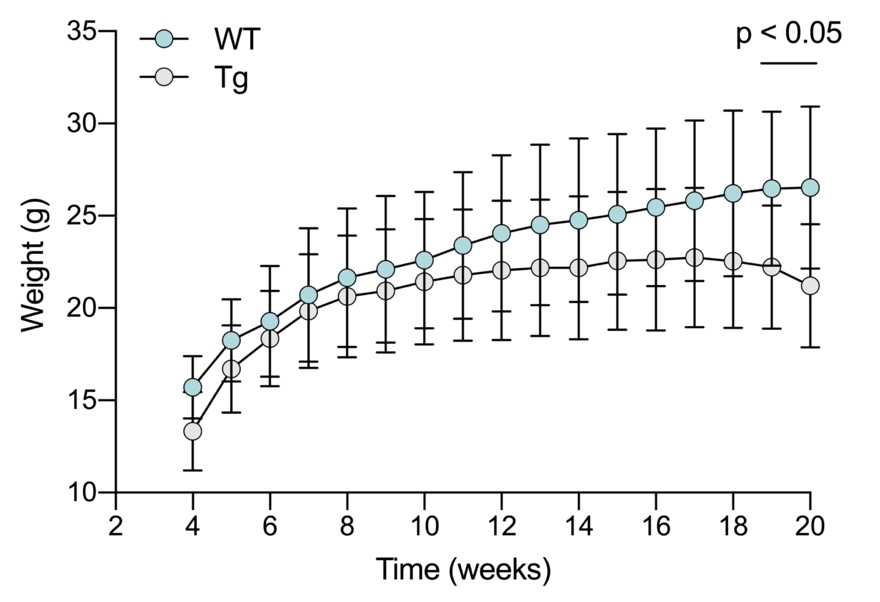
**Suppl. Fig. 1.** Weight evolution across age of wild type and transgenic mice expressed as the average body weight ± SD for each group measured weekly. In comparison to wild-type, the weight of transgenic mice decreases starting from week 19. Data were statistically analyzed by 2-way ANOVA followed by Sidak’s multiple comparison test. A value of p<0.05 was taken as significant.

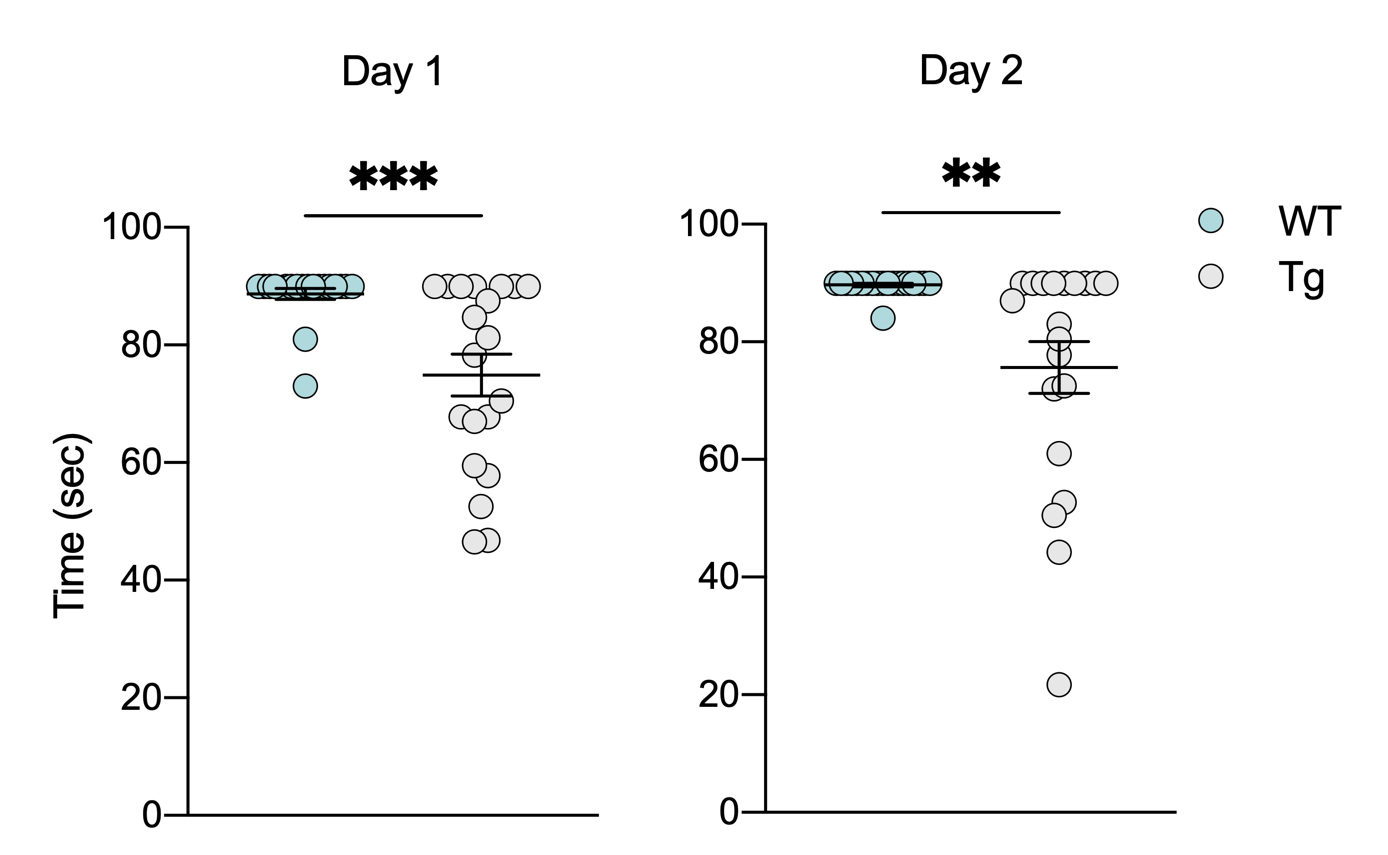

**Suppl. Fig. 2.** Rotarod test performed at week 20 relative to two consecutive days of training at constant speed. Data are expressed as mean of n=20 independent measurements and analyzed by Student t-test, with ** p<0.01 and *** p<0.001.

|  | **Wild type** | **SOD1 G93A** |
| --- | --- | --- |
| Basal | 9.35 ± 1.25 | 7.89 ± 1.87 |
| PMG | 9.85 ± 1.83 | 8.89 ± 1.54 |
| ADP | 44.16 ± 7.00 | 34.91 ± 2.33 |
| S | 72.18 ± 7.92 | 61.44 ± 6.19 |
| CCCP | 106.9 ± 11.39 | 88.17 ± 10.40 |
| Rot | 29.15 ± 5.72 | 23.44 ± 4.73 |
| Ama | 0.76 ± 0.70 | 0.89 ± 0.45 |

**Suppl Tab 1** Raw data relative to oxygen consumption achieved after the addition of substrates, inhibitors and uncouplers in accordance with the SUIT protocol here applied. Basal is the respiration measured after the addition of the samples in the cuvette; PMG, pyruvate, malate and glutamate; S, succinate; Rot, rotenone; AMA, antimycin A. Data are expressed as pmol/s per mg of tissue homogenate and as mean ± SEM of n=10 independent measurements.
